# PqsE acts as an adaptor protein for the quorum-sensing transcription factor RhlR in *Pseudomonas aeruginosa*

**DOI:** 10.1101/2024.12.03.626663

**Authors:** Bilalay V. Tchadi, Jesse J. Derringer, Anna K. Detweiler, Isabelle R. Taylor

## Abstract

*Pseudomonas aeruginosa* is a human pathogen that poses a significant health threat. Pathogenic behaviors of *P. aeruginosa* are under control of the bacterial cell-cell communication system known as quorum sensing (QS). One of the QS master regulators, RhlR, is a receptor/transcription factor that not only relies on binding of its canonical ligand, C4-homoserine lactone (HSL), but additionally requires a protein-protein interaction with the enzyme, PqsE. We constructed heterologous reporter strains in *Escherichia coli* that allow measurements of the reliance of RhlR on C4-HSL and/or PqsE-binding for the ability to activate transcription of three RhlR-regulated genes: *rhlA* (PqsE-independent), *phzM* (PqsE-dependent), and *azeB* (PqsE-suppressed). Analogous assays measuring activation of the three genes in *P. aeruginosa* were performed and the patterns observed correlated tightly with the heterologous reporter assays. These results indicate that binding of PqsE to RhlR is able to fine-tune RhlR transcription factor activity in a promoter-specific manner.

**Importance:** *Pseudomonas aeruginosa* is an opportunistic human pathogen that can cause fatal infections. There exists an urgent need for new, effective antimicrobial agents to combat *P. aeruginosa*. The PqsE-RhlR protein-protein interaction is essential for *P. aeruginosa* to be able to make toxins, form biofilms, and infect host organisms. In this study, we use both non-native models in *E. coli* and measurements of gene expression/toxin production in *P. aeruginosa* to show that the PqsE-RhlR interaction enables fine-tuned gene expression and a heightened ability of *P. aeruginosa* to adapt to external conditions. These findings will be highly valuable as continued efforts are made to design inhibitors of the PqsE-RhlR interaction and test them as potential antimicrobial agents against *P. aeruginosa* infections.

## Introduction

The opportunistic pathogen, *Pseudomonas aeruginosa*, is a Gram-negative bacterium that is responsible for causing life-threatening infections in immuno-compromised individuals. Due to its heightened ability to resist antibiotics, there is a severe lack of treatment options for those who acquire *P. aeruginosa* infections ^1,2^. Resistance to antibiotic treatments as well as several of the virulence phenotypes exhibited by *P. aeruginosa*, such as toxin production and biofilm formation, are under control of the bacterial cell-cell communication process, called quorum sensing (QS) ^3–5^. QS is the process by which bacteria produce, release, and detect small molecule signals (autoinducers) in order to orchestrate coordinated, cell density-dependent group behaviors ^3^. When the extracellular concentration of autoinducers reaches a particular threshold, bacteria make a coordinated lifestyle switch, activating the transcription of genes that promote group behaviors. When *P. aeruginosa* reaches high cell density, genes responsible for producing virulence factors, such as pyocyanin and rhamnolipids, become activated by the QS regulator/transcription factors ^6^.

*P. aeruginosa* has multiple QS circuits in place that regulate a complex network of genes. There are two LuxI/R-type autoinducer synthase/receptor pairs: LasI and LasR, which are responsible for the synthesis and detection of 3-oxo-C12-homoserine lactone (3-oxo-C12-HSL), respectively, and RhlI and RhlR, responsible for the synthesis and detection of C4-homoserine lactone (C4-HSL), respectively ^7,8^. Although *rhlI* expression is under the control of LasR, suggesting that LasR is at the top of the QS regulatory hierarchy, mounting evidence has shown that intact signaling through the Las branch is not required for infection. Rather, mutations in *las* often produce hypervirulent *P. aeruginosa* strains ^9,10^. Contrary to this, RhlR functionality is absolutely required for infection and virulence phenotypes, and to date *rhlR* mutants have not been identified in clinical isolates of *P. aeruginosa* ^11^. It has recently been suggested that understanding of the QS hierarchy be “rewritten”, placing RhlR at the top as the master regulator of quorum-sensing phenotypes in *P. aeruginosa* ^12^.

In addition to a newfound appreciation for the importance of RhlR regulatory function in *P. aeruginosa* virulence, recent progress has been made in understanding how RhlR function differs from that of other LuxR family receptors. In particular, RhlR requires a protein-protein interaction with a metallo-β-hydrolase from a separate QS system, PqsE ^13,14^. When mutations are introduced to either *rhlR* or *pqsE* that disrupt formation of the PqsE:RhlR complex, *P. aeruginosa* can no longer produce pyocyanin or successfully colonize the lungs of a mouse ^15^. It was shown that by forming a complex with RhlR, PqsE enhances the affinity of RhlR for binding its target promoter sequences ^14^. Furthermore, recent structural analyses have illustrated that the two proteins form a functional tetrameric complex, consisting of a PqsE dimer and a RhlR dimer ^16,17^. These recent findings are consistent with the longstanding observation that, unlike other LuxR family receptors, binding of RhlR to its native ligand, C4-HSL, is not sufficient to solubilize the protein when expressed recombinantly in *Escherichia coli* ^18^. This complication had prevented structural characterization of RhlR, prior to the discovery of the PqsE-RhlR protein-protein interaction.

The reliance on a protein-protein interaction for full functionality is a feature that, to date, has only been observed for RhlR of the LuxR family of transcription factors. Recent efforts have focused on understanding what expanded or nuanced activity the interaction with PqsE bestows upon RhlR. A transcriptional analysis revealed RhlR regulates different subsets of genes with varying dependencies on the interaction with PqsE ^19^. Furthermore, a recent chromatin immunoprecipitation and sequencing (ChIP-seq) analysis identified the classes of genes in the RhlR regulon that were dependent on C4-HSL binding, PqsE binding, both, or neither ^20^. Another recent study suggested that PqsE allows for condition-dependent gene regulation ^21^. These recent studies highlight an important feature of RhlR transcription factor activity in that it can be finetuned or adjusted by two independent binding partners: one a protein (PqsE), and the other a small molecule ligand (C4-HSL).

We set out to achieve two goals: 1) characterize regulation of genes representing different classes of RhlR-controlled targets using a heterologous *E. coli* bioluminescent reporter and 2) determine how well the heterologous reporter recapitulates regulation of these genes in *P. aeruginosa*. The *E. coli* system allows for the isolation of PqsE:RhlR:C4-HSL function from other potential regulatory factors present in *P. aeruginosa*, and thus allowed for study of the degree to which C4-HSL and/or PqsE directly determine the functionality of RhlR in a promoter-specific manner. The target gene promoter sequences chosen were those for *rhlA*, the canonical target gene of RhlR responsible for the production of rhamnolipids ^22^, *phzM*, a gene in the phenazine biosynthetic pathway that is necessary for the production of pyocyanin ^23^, and *azeB*, a gene encoding a nonribosomal peptide synthetase that is part of a newly characterized biosynthetic gene cluster responsible for the production of various azetidomonamide and diazetidomonapyridone derivatives with unknown function ^24,25^. These three genes represent different classes of RhlR-regulated genes defined by the previous ChIP-seq analysis with both *rhlA* and *azeB* demonstrating PqsE-independence, and *phzM* depending on both PqsE and C4-HSL ^20^. One intriguing observation from the previous work was that the dependence on either C4-HSL or PqsE was not binary: rather dependence on each binding partner existed on a spectrum with some genes being more or less dependent on C4-HSL (assessed by comparing WT *P. aeruginosa* to the Δ*rhlI* mutant, incapable of producing C4-HSL) or PqsE (assessed by comparing WT *P. aeruginosa* to the Δ*pqsE* mutant). To further investigate this pattern, we probed RhlR transcriptional activity for each specific target promoter in the presence of varying C4-HSL concentrations, as well as when co-expressed with variants of PqsE that have different affinities for RhlR. The PqsE(R243A/R246A/R247A) (from now on referred to as PqsE(NI) for “non-interacting”) variant is completely incapable of self-dimerizing to interact with RhlR, and the PqsE(E182W) variant has a severely weakened interaction with RhlR, which can be circumvented by addition of high concentrations of C4-HSL to restore virulence factor production in strains harboring this mutation. Finally, the PqsE(D73A) variant possesses a similar affinity for RhlR to that of PqsE(WT), but is catalytically inactive. This panel of PqsE mutants as well as the exogenous addition of C4-HSL allowed for the precise measurement of dependence on PqsE versus C4-HSL for full functionality of RhlR.

## Results

### Transcription of rhlA is enhanced by, but not dependent on PqsE binding to RhlR

We were initially interested in characterizing the efficiency of RhlR-activated transcription of its canonical target gene, *rhlA*, when exposed to a range of concentrations of C4-HSL and either in a complex with PqsE or alone. *rhlA* encodes one of the genes necessary for the production of a class of biosurfactants called rhamnolipids. Rhamnolipids play roles in virulence, biofilm formation, and swarming motility ^22,26,27^. Previous characterization of RhlR promoter occupancy identified that binding of RhlR to the *rhlA* promoter was possible in the Δ*pqsE* mutant, but not observed in the Δ*rhlI* mutant. This suggests that RhlR can activate transcription of *rhlA* independently of PqsE, but requires binding of C4-HSL to do so.

Using a previously reported heterologous *E. coli* system to measure transcriptional activation of *rhlA:luxCDABE* via bioluminescence, we determined the sensitivity of RhlR to exogenously added C4-HSL in the presence or absence of an interaction with PqsE (**Figure 1A**). When PqsE was not co-expressed with RhlR in the heterologous system (this strain harbored an empty pACYC184 plasmid with no *pqsE* allele encoded on it), RhlR-activated transcription of *rhlA:luxCDABE* could be fit with a sigmoidal stimulation curve with an EC_50_ of approximately 12 µM C4-HSL. By comparison, the co-expression of PqsE(WT) led to a 100-fold increase in sensitivity of RhlR to C4-HSL (EC_50_ = 118 nM). Three additional PqsE variants were tested in this system, all of which have been previously characterized for their apparent affinity for RhlR ^15^. Briefly, PqsE(D73A) has approximately the same affinity for RhlR as PqsE(WT). PqsE(E182W) is an inhibitor mimetic mutant that has a severely weakened affinity for RhlR and is incapable of driving virulence phenotypes when expressed in *P. aeruginosa*. Finally, PqsE(NI) is completely incapable of forming a complex with RhlR. When tested in the heterologous model for activation of *rhlA:luxCDABE* transcription, the patterns observed for each mutant closely reflected their varying abilities to form a complex with RhlR. Expression of PqsE(D73A) gave a result that closely resembled that of PqsE(WT) (EC_50_ = 205 nM), while expression of PqsE(NI) closely resembled that of no PqsE (EC_50_ = 10.4 µM). Finally, expression of the PqsE(E182W) mutant, which has a weak affinity for RhlR, produced an EC_50_ of approximately 600 nM, reflecting an intermediate level of RhlR sensitivity to C4-HSL.

**Figure 1:**
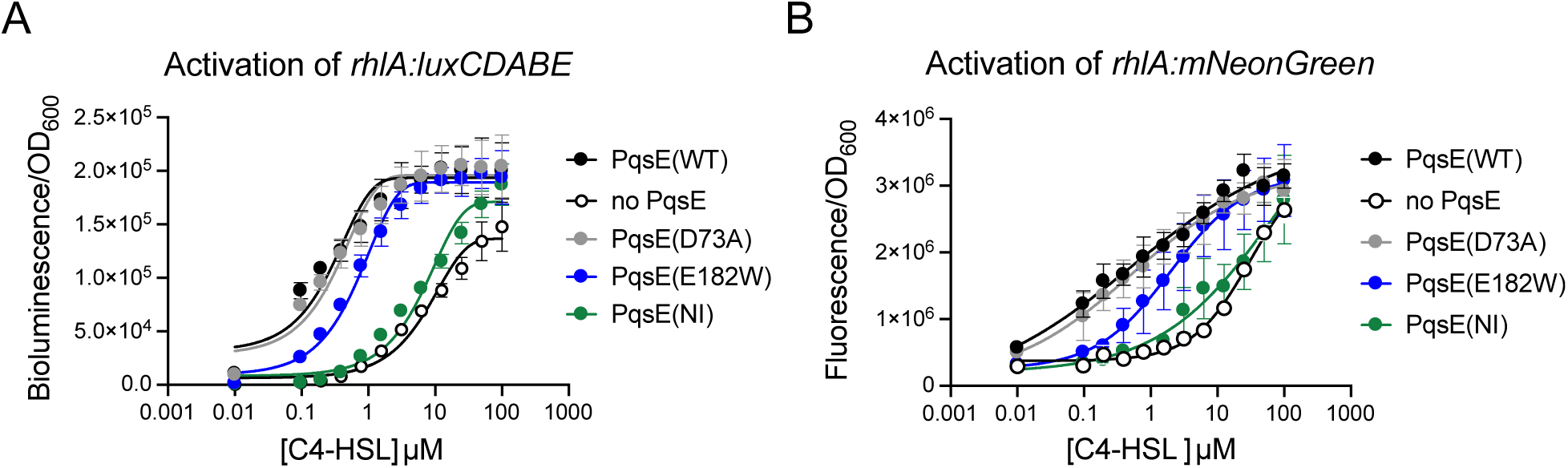
Activation of *rhlA* by RhlR-C4-HSL is enhanced by, but not reliant on PqsE-binding. A) Activation of *rhlA-lux* in *E. coli* is reported as luminescence divided by OD_600_ and the measurements shown were collected after 17 h of growth. B) Activation of *rhlA:mNeongreen* in *P. aeruginosa* is reported as fluorescence (ex: 485 nm em: 535 nm) divided by OD_600_ and measurements shown were collected after 17 h of growth. Stimulation curves were fit to data collected from two independent experiments performed in duplicate. Error bars represent standard deviations.

An analogous experiment was performed in a *ΔrhlI ΔpqsE P. aeruginosa* PA14 strain harboring a *rhlA:mNeonGreen* reporter (**Figure 1B**). Strains expressing each of the *pqsE* alleles tested above on the pUCP18 plasmid were generated in this *rhlA* reporter background in order to measure the dependence of RhlR on C4-HSL binding versus PqsE binding for activating transcription of *rhlA* in *P. aeruginosa*. The resulting C4-HSL stimulation curves closely resembled those generated using the *E. coli* heterologous reporter, albeit with slightly altered concentrations ranges. Both the PqsE(WT) and PqsE(D73A) strains exhibited mid-nanomolar EC_50_ values (516 nM and 498 nM, respectively) and the strains without an intact PqsE-RhlR interaction (the no PqsE and PqsE(NI) strains) were approximately 100 times less sensitive to C4-HSL (EC_50_ values of 31.1 µM and 71.9 µM, respectively). As in the *E. coli* reporter, the *P. aeruginosa* reporter strain harboring the PqsE(E182W) variant displayed an intermediate phenotype, with an EC_50_ of 2.0 µM. The alignment of results in the heterologous reporter with those obtained in *P. aeruginosa* suggest that there are no additional factors present in the native bacterium that influence RhlR transcriptional control over *rhlA* expression other than C4-HSL and PqsE. Rhamnolipid production was quantified for the test strains treated with 0, 0.2, 1, and 10 µM C4-HSL (**Fig S1**). The results, while lacking statistical significance, reflected a similar trend where the dependence on an intact PqsE-RhlR interaction was most apparent at 0 and 0.2 µM C4-HSL. At higher concentrations of C4-HSL, all strains produced similar levels of rhamnolipid to the WT strain.

### Transcription of phzM is dependent on PqsE binding to RhlR and inhibited at high C4-HSL concentrations

The original characterization of the PqsE-RhlR interaction revealed that this interaction was necessary for the ability of *P. aeruginosa* to produce the toxin, pyocyanin. *P. aeruginosa* secretes pyocyanin as a defense mechanism against host or other bacterial cells and pyocyanin production has been correlated to the ability of *P. aeruginosa* to stage an infection ^28,29^. Importantly, pyocyanin is also mildly toxic to *P. aeruginosa* ^30^. The expression of specialized enzymes that decompose the molecule serves as a mechanism to partially protect *P. aeruginosa* from self-toxicity of pyocyanin ^31^. Pyocyanin is a member of a class of molecules called phenazines, produced from the action of the *phzABCDEFGH* operon along with *phzM* and *phzS* ^23^. Accordingly, all of the *phz* genes necessary for pyocyanin production were identified to have strong dependence on both C4-HSL and PqsE for binding of RhlR to their respective promoter regions and transcription ^20^. Notably, *phzM* shares the same promoter region with the *phzA-H1* operon, but is oriented on the opposite strand from the operon. Therefore, it was particularly of interest to determine whether RhlR does in fact regulate *phzM* expression directly.

We adapted the heterologous reporter system to feature the promoter region for the gene, *phzM*, fused to the *luxCDABE* operon and used this reporter to measure activity of RhlR at a range of C4-HSL concentrations and in the presence or absence of the PqsE interaction (**Figure 2A**). In contrast to the *rhlA* reporter, no increase in bioluminescence was observed at any concentration of C4-HSL when PqsE was not co-expressed with RhlR. In the presence of PqsE(WT), RhlR was able to activate transcription producing peak luminescence at approximately 200 nM C4-HSL. Interestingly, at concentrations above ∼500 nM C4-HSL, luminescence began to decrease, falling back down to baseline levels at ∼10 µM C4-HSL. Like with the *rhlA* reporter, the PqsE variants followed a trend that reflects the strength of the interaction with RhlR. The PqsE(D73A) and PqsE(NI) mutants resembled PqsE(WT) and no PqsE, respectively, and expression of the PqsE(E182W) mutant produced an intermediate phenotype.

**Figure 2:**
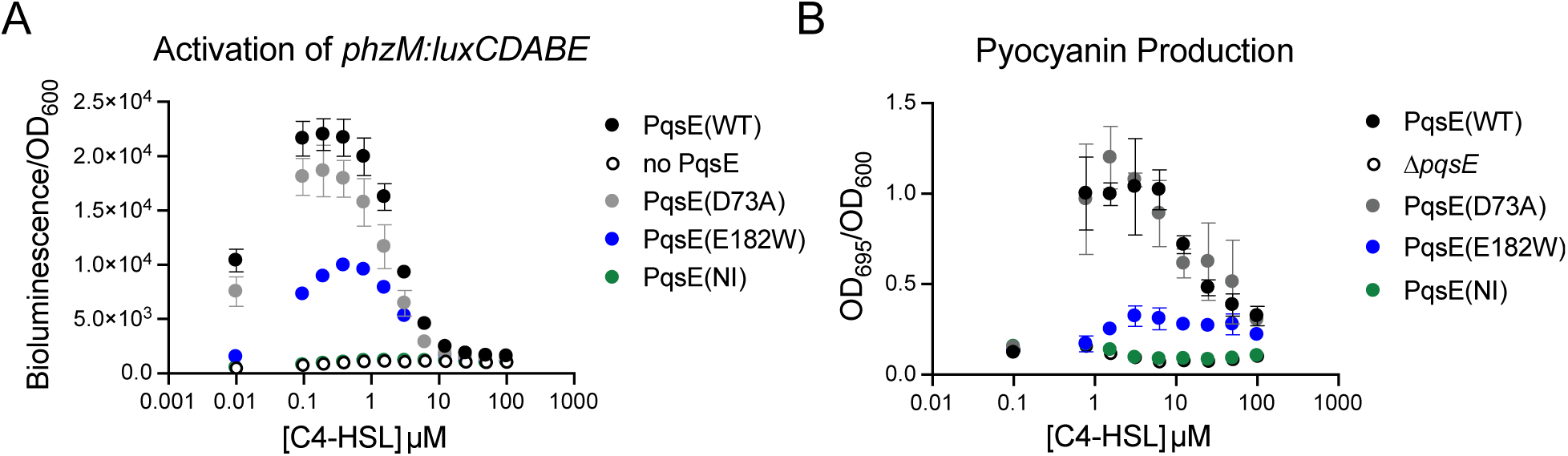
Activation of *phzM* by RhlR-C4-HSL is dependent on PqsE-binding. A) Activation of *phzM-lux* in *E. coli* is reported as luminescence divided by OD_600_ and the measurements shown were collected after 17 h of growth. Results shown are the average of two independent experiments performed in duplicate. B) Pyocyanin production was measured as OD_695_ of the cell-free supernatants and normalized to OD_600_ of the resuspended cell pellets. Measurements were taken after 17 h of growth. Results shown are the average of two biological replicates performed in duplicate. Error bars represent standard deviations.

The enzyme, PhzM, catalyzes the methylation of phenazine-1-carboxylic acid, a necessary step in the production of pyocyanin. Therefore, in order to determine how well results from the heterologous reporter replicated transcriptional regulation of *phzM* in *P. aeruginosa*, we measured pyocyanin production in Δ*rhlI* Δ*pqsE P. aeruginosa* strains expressing the panel of PqsE variants on the pUCP18 plasmid with exogenous addition of C4-HSL at a range of concentrations (**Figure 2B**). As in the heterologous reporter, the strains that lacked an intact PqsE-RhlR interaction (no PqsE and PqsE(NI)) did not produce pyocyanin at any concentration of C4-HSL. Also mirroring the heterologous reporter, both PqsE(WT) and PqsE(D73A) were able to produce pyocyanin, with peak production observed at approximately 1 µM C4-HSL. As with the heterologous reporter, pyocyanin production decreased to near baseline levels at higher concentrations of C4-HSL, consistent with previous observations ^32^. And finally, the PqsE(E182W) mutant, which has a weakened affinity for RhlR, displayed an intermediate phenotype, where peak pyocyanin production amounted to 26% of that produced by the strain expressing PqsE(WT). This result correlates well with previous studies in which PqsE(E182W)-expressing strains exhibited a similar impairment in pyocyanin production when *rhlI* was not deleted.

### Transcription of azeB is enhanced by PqsE binding in the absence of C4-HSL, but suppressed by binding of PqsE to RhlR:C4-HSL

The *aze* operon consists of 10 genes encoding enzymes that collectively produce a series of azetidine-containing natural products, including the molecule azabicyclene. This gene cluster is highly conserved among *P. aeruginosa* strains, with further conservation among 12 additional *Pseudomonas* species ^24^. While the precise roles of these natural products is not known, deletion of certain genes in the cluster were shown to increase the ability of *P. aeruginosa* PAO1 to colonize the mouse lung ^33^. Interestingly, the entire *aze* operon, along with *azeA*, have been identified as being part of the core RhlR regulon in clinical isolates of *P. aeruginosa* harboring mutations in the *las* system ^34^. Furthermore, binding of RhlR to the *azeB* promoter was shown to be independent of PqsE ^20^. For all of the above reasons, regulation of the *az*e gene cluster by PqsE:RhlR was of great interest. We therefore constructed luminescent reporters to study transcription of *azeB* in both *E. coli* and *P. aeruginosa*.

In *E. coli*, we found that the dependence of RhlR on C4-HSL and PqsE binding followed a unique pattern that had not been observed for any genes of interest previously. In the absence of C4-HSL, strains harboring an intact PqsE:RhlR complex (PqsE(WT) and PqsE(D73A)) produced greater signal than those in which the PqsE-RhlR interaction was abolished (no PqsE and PqsE(NI)), with the strain expressing PqsE(E182W) producing an intermediate level of luminescence. This trend became completely reversed with the addition of C4-HSL, where luminescence decreased in the strains expressing PqsE(WT) and PqsE(D73A) and increased in those lacking a stable interaction between PqsE and RhlR. Peak activity of RhlR was reached at ∼40 nM C4-HSL in the strain expressing PqsE(E182W) and 100 nM in both the strain expressing no PqsE and the strain expressing PqsE(NI). High concentrations of C4-HSL decreased luminescence, bringing the signal down to baseline levels by 10 µM C4-HSL.

In *P. aeruginosa*, activation of *azeB:luxCDABE* followed a similar trend to that observed in *E. coli*, at slightly altered concentration ranges of C4-HSL. As in *E. coli*, in the absence of C4-HSL, PqsE(WT) and PqsE(D73A) were both able to slightly boost activation of the transcriptional reporter. With the addition of mid-nanomolar concentrations of C4-HSL, the trend was reversed, showing that strains in which the PqsE-RhlR interaction was intact produced lower luminescence values. The signal reached peak levels at ∼400 nM C4-HSL in the strain expressing PqsE(E182W), and ∼1 µM C4-HSL in both the strains expressing no PqsE and PqsE(NI). Higher concentrations of C4-HSL decreased activation of *azeB:luxCDABE* and reached near baseline levels at ∼100 µM C4-HSL.

**Figure 3:**
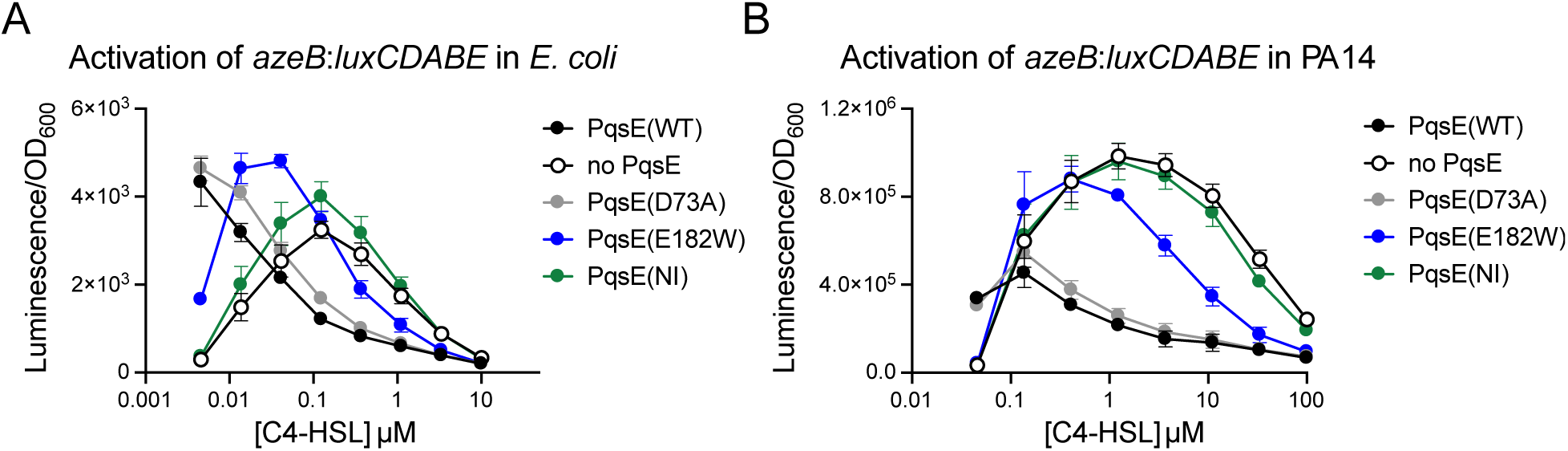
Activation of *azeB* by RhlR:C4-HSL is suppressed by PqsE-binding. A) Activation of *azeB-lux* in *E. coli* is reported as luminescence divided by OD_600_ and the measurements shown were collected after 17 h of growth. B) Activation of *azeB-lux* in *P. aeruginosa* is reported as luminescence divided by OD_600_ and measurements shown were collected after 17 h of growth. Results shown are the average of two independent experiments performed in duplicate. Error bars represent standard deviations.

## Discussion

In this study, we used a heterologous reporter system, along with a well-characterized set of PqsE variants with varying affinities for the transcription factor, RhlR, to precisely determine contributions of each of RhlR’s binding partners, PqsE and C4-HSL, to the regulation of genes within the RhlR regulon. In each of the three cases explored here, expression patterns observed for RhlR target genes in the heterologous *E. coli* model closely resembled the patterns observed for these genes and their virulence factor products in *P. aeruginosa*. This finding suggests that there are no other regulatory factors in *P. aeruginosa* other than PqsE and C4-HSL that influence RhlR-dependent activation of the genes *rhlA*, *phzM*, and *azeB*. It is worth noting that the effect of PqsE-binding on RhlR activity differed depending on the identity of the promoter region of interest. It was also notable that high concentrations of C4-HSL had a stimulatory effect in the case of *rhlA* transcription, but an inhibitory effect in the cases of both *phzM* and *azeB*. Because these expression patterns were observed in both *P. aeruginosa* and a non-native bacterium (*E. coli*), we suggest that this promoter-specific activity arises from the existence of multiple active conformations of RhlR, possibly involving multimeric species of the transcription factor. Further work will be required to determine if this is the case.

The *P. aeruginosa* QS regulator/transcription factor, RhlR, is a unique LuxR family receptor. Not only does it rely on binding of its native ligand, C4-HSL, but it additionally requires an interaction with the protein, PqsE, for full functionality. Although eukaryotic transcription factors are known to rely on many complex protein-protein interactions ^35^, in the realm of prokaryotic gene regulation, this is unusual. It must be noted, however, that the binding of PqsE is not necessary for RhlR’s ability to activate transcription of some genes in its regulon (*rhlA* being the example in this study). Rather, it appears that RhlR may be able to adopt multiple conformational states, one of which is stabilized by PqsE. The PqsE-stabilized state seems to be the only state capable of binding some promoters, such as for *phzM*, but not the only state capable of binding other promoters, like that of *rhlA*. And in this study, it was noted that transcription of another subset of RhlR-regulated genes, such as *azeB*, is suppressed by binding of PqsE to RhlR. The expression patterns observed for *azeB* were complicated, as it appeared PqsE-binding does activate RhlR in the absence of C4-HSL, but even at low nanomolar concentrations of C4-HSL, PqsE-binding switched to suppress activation of this gene. Taken together, we propose that PqsE is acting as an adaptor protein for RhlR, adjusting its transcription factor activity in a promoter-specific manner, and in response to varying concentrations of C4-HSL.

It is easy to imagine why *P. aeruginosa* would have evolved such a mechanism to finetune regulation of genes responsible for the production of its various virulence factors. Through QS, *P. aeruginosa* senses the extracellular concentration of C4-HSL in order to gauge the number of neighboring *P. aeruginosa* cells in its vicinity. The production of rhamnolipids confers several benefits to *P. aeruginosa* in addition to toxicity against other species of bacteria, including biofilm formation, swarming motility, and nutrient acquisition ^36^. There is no drawback to the production of rhamnolipids when *P. aeruginosa* is at high cell density. Pyocyanin acts as a toxin against other competing species of bacteria, as well as host cells. However, pyocyanin is also toxic to *P. aeruginosa* itself ^30,31^. Therefore it is logical that, when exposed to high concentrations of C4-HSL (signifying many neighboring *P. aeruginosa* cells) *P. aeruginosa* should stop expressing the genes involved in pyocyanin production. It is intriguing, therefore, to observe that PqsE suppresses RhlR-activated transcription of some genes, such as *azeB*. It begs the question of what unintended effects might arise from the inhibition of the PqsE-RhlR interaction, which is currently being explored as an antibiotic route. More work must be done to characterize the roles of molecules produced by the *aze* biosynthetic pathway. Regardless, the tunability of the PqsE-RhlR system serves as an example of the heightened adaptability of *P. aeruginosa*, and most likely contributes to pathogenicity of this notorious bacterium.

## Materials and Methods

### Strains, media, and molecular procedures

All *P. aeruginosa* strains were generated in the UCBPP-PA14 parental strain background and all *E. coli* reporter strains were generated in the Top10 parental strain background. Unless otherwise stated, all strains were grown in Luria-Bertani broth and supplemented with appropriate selective antibiotics at the following concentrations: carbenicillin (400 µg/mL), ampicillin (200 µg/mL), kanamycin (100 µg/mL), and chloramphenicol (10 µg/mL). Plasmids were transformed into *P. aeruginosa* strains as described ^37^ and into *E. coli* strains by electroporation. A list of all strains featured in this study is included in the Supporting Information (**Table S1**).

### Generation of azeB-lux reporters

Both the pCS26-*azeB-luxCDABE* plasmid (*E. coli* reporter) and a pUC18T-mini-Tn7T-P*azeB-lux*-Tp plasmid for construction of a *P. aeruginosa azeB* reporter were assembled following a previously established protocol ^38^. Briefly, the 470 bp region upstream of *azeB* and each vector were amplified to introduce ∼20 bp overlaps. The amplified vectors were digested with Dpn1 and 1 µL of each (vector and promoter region insert) were transformed into ultracompetent XL-10 Gold cells (Agilent) for *in vivo* assembly. Positive clones were confirmed by sequencing, and the isolated pCS26-*azeB-luxCDABE* plasmid was transformed into *E. coli* Top10 to generate the heterologous reporter. The pUC18T-mini-Tn7T-P*azeB-lux*-Tp plasmid was transformed into Top10 and mated with *P. aeruginosa* PA14 to stably integrate the *azeB-luxCDABE* fusion at the att-Tn7 site next to *glmS*.

### Heterologous reporters of RhlR transcription factor activity

*E. coli* strains harboring plasmids with *rhlR* driven by the P_BAD_ promoter, *luxCDABE* under the P*rhlA*, P*phzM*, or P*azeB* promoter on the pCS26 vector, and either the pACYC184 vector or pACYC184 harboring WT *pqsE* or mutant *pqsE* alleles were grown with shaking overnight at 37 °C in LB medium supplemented with ampicillin, kanamycin, and chloramphenicol. Overnight cultures were diluted 1:1000 into fresh LB with antibiotics, 0.1% arabinose, and varying concentrations of C4-HSL then added to the wells of a white, clear-bottomed 96-well plate (Corning 3903) to a total volume of 100 µL per well. Plates were incubated in the plate-reader (Molecular Devices iD5) at 30 °C with periodic shaking for 24 h and luminescence and OD_600_ were measured every 10 minutes. Bioluminescence values are reported as the luminescence measurement divided by OD_600_ and the timepoints shown are the time at which peak signal was observed (typically at 17 h).

### rhlA:mNeonGreen *assay*

*ΔrhlI ΔpqsE P. aeruginosa* PA14 strains harboring *mNeonGreen* under the *rhlA* promoter, and either a pUCP18 control vector or pUCP18 harboring WT *pqsE* or mutant *pqsE* alleles were grown with shaking overnight at 37 °C in LB medium supplemented with carbenicillin. Overnight cultures were diluted 1:1000 into fresh LB with carbenicillin and varying concentrations of C4-HSL then added to the wells of a round-bottomed 96-well plate (Corning 3797) to a total volume of 100 µL per well. The lid and a Breathe-EASIER plate sealing film (Diversified Biotech BERM-2000) were applied to minimize evaporation during shaking overnight at 37 °C. The following day, cells were pelleted by centrifugation at 2000 rpm for 20 min at 4 °C. The supernatant was removed and cell pellets were resuspended in 100 µL of PBS (137 mM NaCl, 2.7 mM KCl, 10 mM Na_2_HPO_4_ 1.8 mM KH_2_PO_4_, pH 7.4), per well. Cell suspensions were transferred into the wells of a black, clear-bottomed 96-pell plate (Corning 3904) and fluorescence (excitation at 485 nm; emission at 535 nm) and OD_600_ were measured in a plate-reader (Molecular Devices iD5).

### Rhamnolipid quantification

Cultures of *P. aeruginosa* strains were grown from single colonies in LB medium at 37 °C with shaking overnight. Cultures were diluted 1:1000 in fresh LB supplemented with antibiotics and C4-HSL at the specified concentrations (0.1% DMSO), and grown with shaking at 37 °C for 20 h. Cultures were then pelleted by centrifugation at 12,000 rpm for 10 min at 25 °C and the cell-free supernatants were subsequently passed through a 0.22 µm filter. The supernatant was then acidified to pH 2 using HCl and incubated at room temperature overnight. Samples were clarified by centrifugation at 4,000 rpm for 20 min at 25 °C, then decanted and extracted twice with 1 mL chloroform:methanol (2:1) ^39^. The organic fractions were dried using a rotary evaporator, then resuspended in a 53% H_2_SO_4_ solution with 0.19% orcinol (w/v) ^40^. Samples were incubated at 80 °C for 30 min, then cooled at room temperature for 15 min. Absorbance was measured at 421 nm in a plate-reader (Molecular Devices iD5) and compared to rhamnose standards.

### Pyocyanin assay

Overnight cultures of Δ*rhlI* Δ*pqsE P. aeruginosa* strains harboring the empty pUCP18 vector or the vector expressing WT *pqsE* or mutant *pqsE* alleles were grown from single colonies in LB medium at 37 °C. Cultures were diluted 1:1000 in 2 mL of fresh LB liquid medium supplemented with carbenicillin and C4-HSL (1% DMSO) at the specified concentrations and grown with shaking at 37 °C for 17 h. Cells were then pelleted by centrifugation and OD_695_ of the clarified supernatants was measured in a Genesys 20 ThermoSpectronic spectrophotometer. Pellets were resuspended in PBS and OD_600_ was measured to determine cell density of each sample. Pyocyanin production is reported as the OD_695_ of clarified supernatant normalized to the OD_600_ of the resuspended pellet.

### P. aeruginosa azeB:lux *assay*

*ΔrhlI ΔpqsE P. aeruginosa* PA14 strains harboring *luxCDABE* under the P*azeB* promoter, and either a control pUCP18 vector or pUCP18 harboring WT *pqsE* or mutant *pqsE* alleles were grown with shaking overnight at 37 °C in LB medium supplemented with carbenicillin. Overnight cultures were diluted 1:1000 into fresh LB with carbenicillin and varying concentrations of C4-HSL then added to the wells of a white, clear-bottomed 96-well plate (Corning 3903) to a total volume of 100 µL per well. Plates were incubated in the plate-reader (Molecular Devices iD5) at 30 °C with periodic shaking for 17 h and luminescence and OD_600_ were measured every 10 minutes.

## Supporting Information

Results of rhamnolipid quantification experiments and a list of all strains used in this study (PDF).

## Acknowledgments

We thank members of the Taylor Lab for helpful advice and discussions. We also thank Dr. Jon Paczkowski for cloning advice and providing feedback on the manuscript.

## Author Contributions

B.V.T., J.J.D., and A.K.D. conducted experiments. B.V.T., J.J.D., A.K.D., and I.R.T. designed experiments and prepared the manuscript.

